# Unveiling Lipid Dysregulation: Lipidomics of Mouse Brain and Isolated Myelin in Niemann–Pick Disease Type C1

**DOI:** 10.64898/2026.02.07.704575

**Authors:** Koralege C. Pathmasiri, Stephanie M. Cologna

## Abstract

Niemann–Pick Disease Type C1 (NPC1) is a fatal, neurodegenerative disorder, characterized by lysosomal lipid accumulation and dysmyelination. Previous studies have documented some lipid abnormalities in the null mouse (*Npc1^−/−^)* focused on the whole brain and liver. However, the specific lipidomic alterations in severely affected brain regions, such as cerebellum and isolated myelin remain understudied. We present a comprehensive LC-MS-based lipidomic analysis of the cerebellum and cortex of *Npc1^−/−^* mice during disease progression stages, along with the first comprehensive characterization of the myelin lipidome in NPC1 disease. Our results reveal that the cerebellum accumulates lipid species, including sphingolipids and glycerophospholipids progressively, while the cortex shows an overall decline in lipid levels, indicating region-specific lipid dysregulation. Notably, bis(monoacylglycero)phosphates and their precursors—including lysophosphatidylglycerol and hemibismonoacylglycerophosphate exhibit significant accumulation, with a preference for docosahexaenoic acid (DHA)-containing species. Despite known cholesterol storage defects in NPC1, we observed reduced free cholesterol levels in both regions, which we attribute to myelin loss. Myelin-specific lipidomics demonstrated extensive dysregulation, particularly in cortical myelin, including severe losses in sulfatides, ether-lipids, and acylcarnitine, alongside striking accumulation of hydroxy-ceramides. These findings identify novel lipid alterations in brain subregions and myelin, offering critical insight into the lipid perturbations under the loss of NPC1, and highlight lipid targets that may be crucial for therapeutic intervention and biomarker development.

## Introduction

Niemann–Pick Disease Type C1 (NPC1) is a rare, autosomal recessive lipid storage disorder marked by defective intracellular transport of cholesterol and other lipids, such as sphingolipids, leading to their accumulation in late endosomes and lysosomes^1^. The disease affects multiple organs, including the central nervous system, liver, and spleen. Progressive neurodegeneration is a hallmark of NPC1 disease, with individuals exhibiting symptoms such as ataxia, cognitive decline, vertical supranuclear gaze, seizures, and dementia^2^. The neurological impairment reflects widespread disruption of neuronal function and viability, particularly in the cerebellum, but also in other brain regions. A characteristic neurological feature in the disease is the loss of cerebellar Purkinje cells, which are large neurons critical for motor coordination and balance^3,4^. Purkinje cell degeneration in NPC1 is progressive and regionally patterned, progressing in a rostral to caudal manner, correlating with worsening motor deficits such as tremors and impaired coordination^5^. Given the importance of lipids as structural and signaling biomolecules, a comprehensive view of altered lipids beyond cholesterol and sphingolipids is crucial for future therapeutic development and evaluation.

The impaired intracellular lipid trafficking in NPC1 disease leads to the accumulation of unesterified cholesterol and sphingolipids in late endosomes and lysosomes, creating a lipid imbalance within the cell^6^. These lipid imbalances disrupt cellular homeostasis, particularly in the central nervous system, contributing to progressive neurodegeneration. Investigating the lipidome alterations associated with NPC1 can reveal disease-specific biomarkers, provide insights into disease pathogenesis, and inform the development of targeted therapeutic strategies aimed at restoring lipid balance.

In addition to progressive neurodegeneration, dysmyelination – defined as improper or incomplete myelin formation-has been reported but is also understudied in NPC1 disease^7,8^. Emerging evidence suggests that defects in myelin formation and maintenance also may contribute to neurological symptoms in NPC1 disease^7,8^. Myelin, which is essential for rapid signal conduction and axonal integrity, is disrupted in both NPC1 individuals and animal models of NPC1, as evidenced by reduced expression of myelin-associated proteins, altered oligodendrocyte function, and structural abnormalities in the white matter tract^7,9^. The accumulation of lipids within oligodendrocytes and their precursor cells likely impairs the normal myelination process, compounding the effect of neuronal degeneration^10^.

NPC1 disease research has benefited from the development and use of multiple animal models, including several mouse models^11,12^, a feline model^13^, and a zebrafish model^14^. Among these, mouse models are the most widely used due to their genetic tractability, well-characterized phenotype, and suitability for both mechanistic studies and therapeutic development^15^. In particular, the *Npc1* null (*Npc1^−/−^*) mouse model, which harbors a targeted disruption of the *Npc1* gene, has become a foundational tool in the field^15,16^. Although complete loss-of-function mutations are rarely observed in human NPC1 patients, this model exhibits hallmark features of the human disease, including intracellular accumulation of unesterified cholesterol and glycosphingolipids, progressive neurodegeneration, motor deficits, and hepatosplenomegaly^17^.

In the current study, we utilized the *Npc1* null mouse model at three defined stages of disease progression: 3 weeks (pre-symptomatic), 7 weeks (symptomatic), and 9 weeks (late-stage/severe phenotype). These time points are critical for understanding the temporal sequence of pathophysiological events associated with NPC1 deficiency, ranging from early molecular alterations to clear neurological impairment. By analyzing the lipidome across the disease progression, we reveal crucial alterations in the brain lipidome that accompany the advancement of the pathological condition. Towards that end, we have focused this study on the cerebellum, cortex, and enriched myelin lipidome of these regions in the *Npc1* null mouse brain.

## Results

### Overall lipid perturbations in *Npc1^−/−^* mice cerebellum and cortex across disease progression

The sphingolipid and free fatty acid profiles of the *Npc1^−/−^* mice model have been reported in previous studies, with a focus on the whole brain^18^ and liver^19,20^. In this study, we used an LC-MS lipidomic approach to expand the scope of the lipidome investigation and have characterized more than 30 lipid classes (**Figure S1A, S1B**). Since the cerebellum is more severely affected in NPC1 disease compared to other brain regions^21^, we have dissected the mouse brain into cerebellum and cortex to explore the lipid changes in these areas separately. Our data demonstrate that the lipid classes present in the cerebellum and cortex are similar, as are the numbers of lipid species identified within each class. However, the cerebellar lipidome of *Npc1^−/−^* mice exhibits a continued increase in the number of accumulated lipid species with disease progression, while the cortex lipidome has more downregulated lipids with increased disease burden (**Figure 1**). With respect to the total number of dysregulated lipids, the cortex exhibits a larger number of differential lipids at each time point. The cortex has 182, 219, and 230 differential lipids at 3,7, and 9 weeks, while the cerebellum has 162, 112, and 144 differential lipids at 3, 7, and 9 weeks, respectively (**Figure 1**). The results of unbiased hierarchical clustering analysis and principal component analysis demonstrate distinct separation between the *Npc1^−/−^*and *Npc1^+/+^* samples, highlighting significant alterations in the lipidome throughout each time point of the disease progression (**Figures S2 and S3**). To assess how known lipid changes are represented in the cortex and cerebellum lipidomes individually, and to uncover novel alterations, each lipid class within each lipid category was further examined.

**Figure 1.**
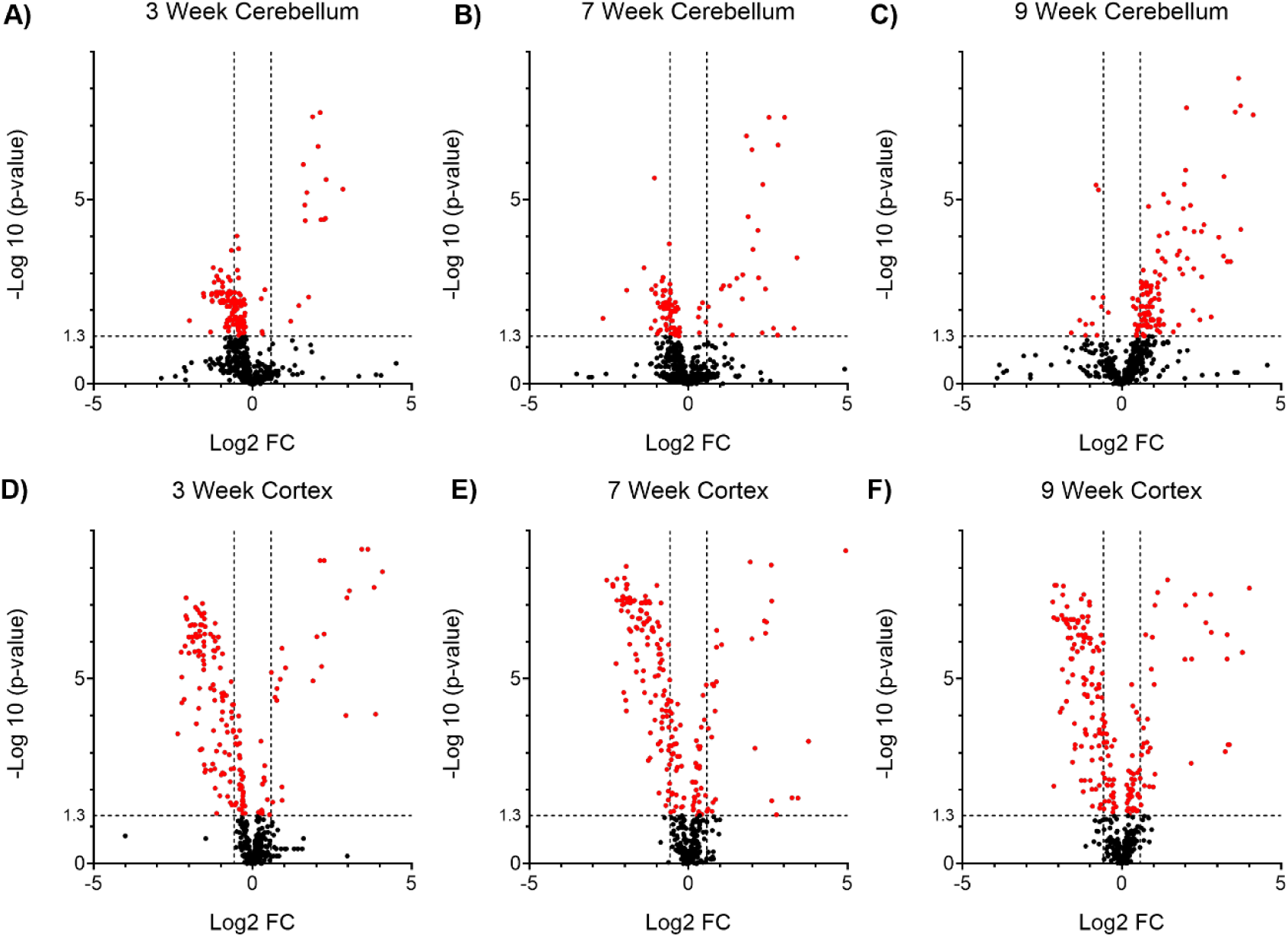
Volcano plots for the differential LC-MS lipidomics in the NPC1 mouse cerebellum and cortex. An unpaired t-test was performed between the *Npc1^−/−^* and *Npc1^+/+^* at each time point, and the Log2 Fold change (FC) was plotted against the -Log 10 p-value to generate volcano plots. A) 3 weeks cerebellum B) 7 weeks cerebellum C) 9 weeks cerebellum D) 3 weeks cortex E) 7 weeks cortex F) 9 weeks cortex. All the data points with p-value < 0.05 are highlighted in red. Dotted lines are added to intercept the x-axis at FC = 1.5 and FC = −1.5 to show the FC cut off and for the y-axis at p-value = 0.05.

### Glycerophospholipid alterations in *Npc1^−/−^* mouse cerebellum and cortex

Previous studies have shown the bis(monoacylglycero)phosphate (BMP) lipid, also known as lysobisphosphatidic acid (LBPA), accumulates in NPC1-deficient cells^22,23^. These BMP lipids are known to be enriched in the intraluminal vesicles (ILVs) located in late endosomal/lysosomal (LE/LY) compartments^24,25^. It has also been reported that the BMP lipids may bind to the NPC2 cholesterol transporter and stimulate the cholesterol transportation^23^. Moreover, BMP lipids are negatively charged at low pH environments such as the lysosome (pH 4.5–5) and are suggested to facilitate the binding of acid hydrolase enzymes to intraluminal vesicle (ILV) membranes, thereby enabling lysosomal lipid degradation/metabolism^26,27^. Our data shows that the BMP lipids are the major glycerophospholipids that exhibit the highest fold accumulation in both the cerebellum and the cortex (**Figure 2A, 2B**). While the BMP precursor, Lyso-phosphatidylglycerol (LPG) can be highlighted as the most prominent lyso glycerophospholipid alteration in both brain regions (**Figure 2C, 2D**). In addition, BMPs are one of the few glycerophospholipids that exhibit accumulation at the 3-week time point in both the cerebellum and cortex. It is also important to note that all the identified BMP lipid species, regardless of fatty acyl chain composition, show significant accumulation at all time points in both brain regions. The fatty acid composition of these BMP lipids reveals that the major fatty acid incorporated into these lipids is docosahexaenoic acid (DHA), which is an omega-3 fatty acid essential in brain development^28^ and neuronal function^29^. In the cerebellum, out of the 13 BMP species identified, 8 of them contain DHA, while in the cortex, 5 of the eight identified BMP lipids contain DHA (**Figure S4**). Other polyunsaturated fatty acids incorporated into BMP lipids include arachidonic acid, docosatetraenoic acid, docosapentaenoic acid, linoleic acid, and eicosapentaenoic acid. While these polyunsaturated fatty acids account for more than 70% of the BMP fatty acids in both brain regions, the major monounsaturated fatty acid incorporated into BMP is found to be oleic acid.

**Figure 2.**
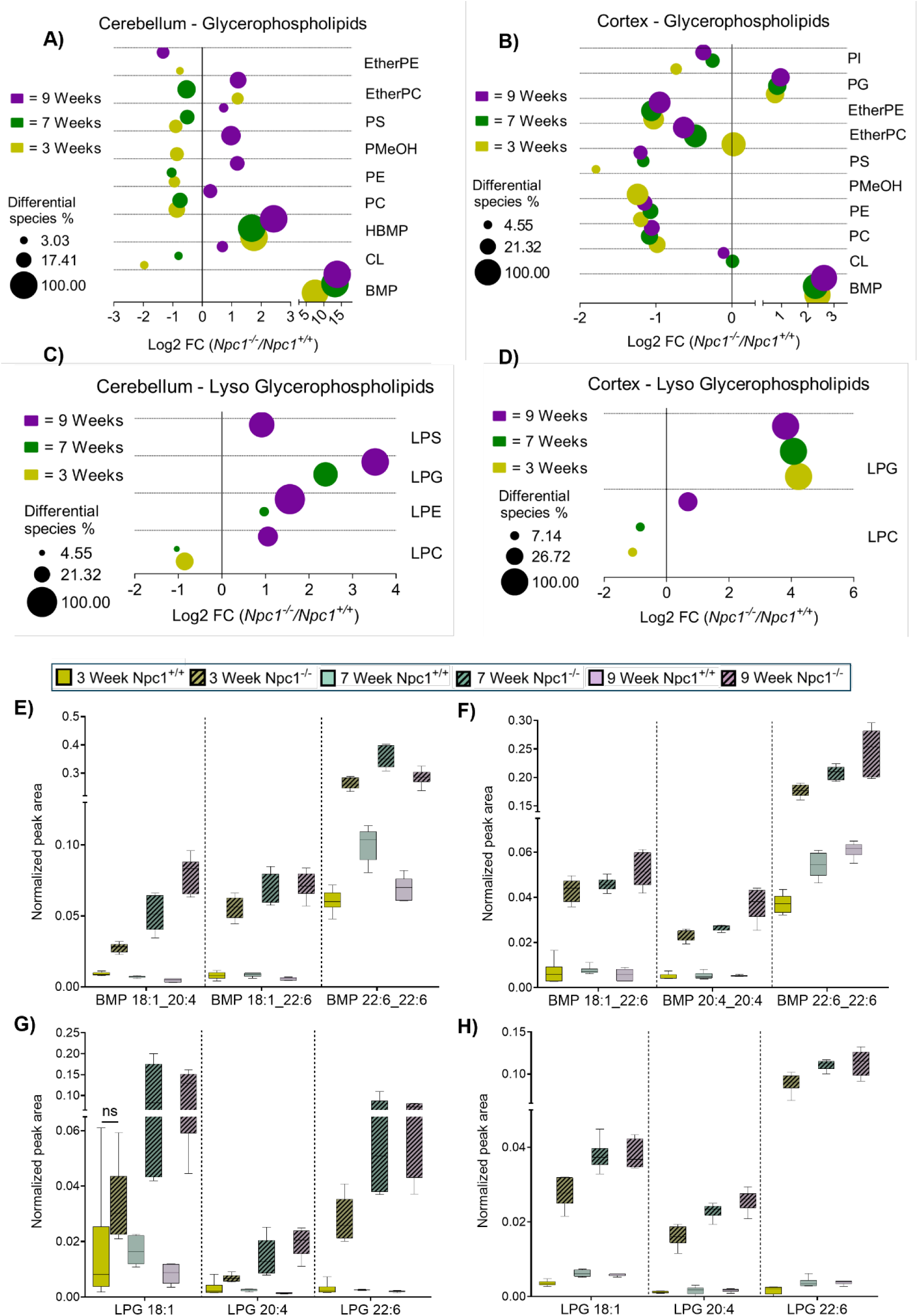
Glycerophospholipids and Lyso Glycerophospholipids show statistically significant changes in the *Npc1^−/−^* tissues. A) Cerebellum and B) Cortex differential Glycerophospholipids at each time point. C) Cerebellum and D) Cortex differential Lyso-Glycerophospholipids at each time point. Summed, normalized abundances of the differential lipids were used to calculate the Log2 Fold change (FC) and used to create the bubble plots. Bubble size is scaled to represent the percentage of significantly different lipid species compared to number of total identified lipid species in each lipid class. Unpaired t-tests were conducted between *Npc1^−/−^* and *Npc1^+/+^* for each time point and each tissue to calculate the p-values and the lipid changes with a p-value < 0.05 and FC > 1.5 were considered significant. All the lipids included in the calculations show statistically significant differences between the *Npc1^−/−^* and *Npc1^+/+^* groups. The three most prominent Bis(monoacylglycero)phosphate (BMP) species show severe accumulation in the *Npc1^−/−^* E) cerebellum F) cortex tissues. G) cerebellum H) cortex Lysophosphatidylglycerol (LPG) species that show significant accumulation in the null. Peak areas normalized to internal standards were used to create the box and whisker plots.

Three of the highest-abundant BMP species found in the cerebellum are BMP 18:1_20:4, BMP 18:1_22:6, and BMP 22:6_22:6, while in the cortex, BMP 18:1_22:6, BMP 20:4_20:4, and BMP 22:6_22:6 are the highest-abundant BMPs (**Figure 2E, 2F**). Interestingly, the corresponding precursor lyso-phosphatidylglycerol (LPG) species also shows pronounced accumulation in both brain regions (**Figure 2G, 2H**). This finding expands the current knowledge of BMP accumulation in NPC1 disease. Consistent with the BMP profile, all LPG species show marked accumulation during disease progression, except for mono-unsaturated LPG 18:1 in the cerebellum at the 3-week time point. The most pronounced accumulation in both brain regions is DHA-containing LPG species, mirroring the trends observed in BMP species. In the process of BMP synthesis, LPG is formed by the hydrolysis of PG by the phospholipase A (PLA) enzymes. Primarily, the lysosomal PLA2 has been identified as the enzyme responsible for initiating BMP synthesis by converting PG into LPG^30^. This resulting LPG then gets acylated to synthesize BMP, and this reaction from LPG to BMP is shown to be catalyzed by phospholipases D3 and D4 (PLD3 and PLD4) and Bis(monoacylglycero)phosphate synthase CLN5 enzymes^31^.

### Sphingolipid alterations in *Npc1^−/−^* mouse cerebellum and cortex

Apart from synthesis through LPG intermediate, another potential BMP synthesis pathway is via hemi-BMP (HBMP) intermediate^32^. Hemi-BMP, also known as acyl-PG, is an uncommon acyl glycerol phospholipid that consists of three fatty acyl chains^33^. Consistent with the accumulation of LPG and BMP discussed above, our results indicate a substantial increase in HBMP levels in *Npc1^−/−^* cerebellum and cortex at all 3 time points (**Figure 3A, 3B**). Additionally, the fatty acid composition of the HBMP with elevated levels reveals the dominance of DHA incorporation. These findings suggest that both BMP synthesis pathways discussed above contribute to the BMP accumulation observed in the *Npc1^−/−^*. While HBMP is suggested to be an intermediate in BMP biosynthesis, it is also important to note that some reports indicate HBMP may instead be an acylation product of BMP^33^.

**Figure 3.**
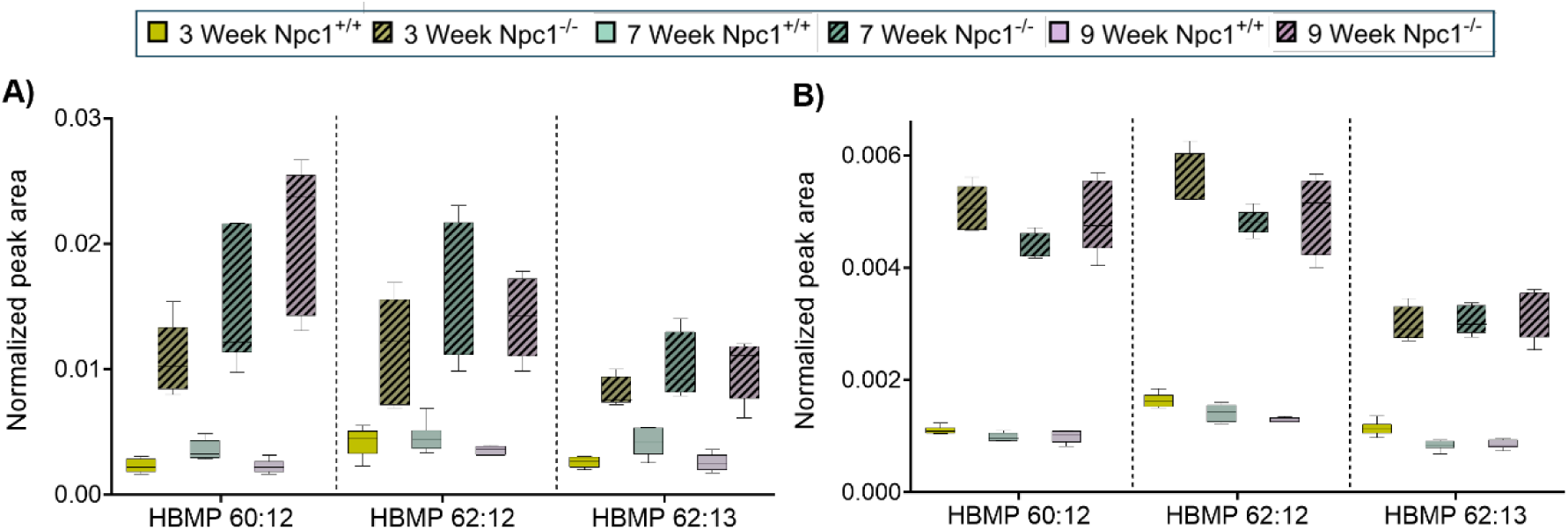
Hemibismonoacylglycerophosphate (HBMP) lipids that show significant accumulation in the cerebellum and cortex tissues at all 3-time points A) Cerebellum and B) Cortex. Peak areas normalized to internal standards were used to create the box and whisker plots.

Another lipid category that is severely affected in NPC1 disease is sphingolipids. In this study, we report multiple sphingolipid classes, including ceramides and ganglioside accumulations, as well as a novel finding of psychosine accumulation in the *Npc1^−/−^* mice brain. Moreover, we point out some sphingolipids that deviate from the general trend of sphingolipid accumulation in NPC1 disease. The sphingolipids exhibiting decreased levels in the *Npc1^−/−^* cerebellum include non-hydroxy ceramides (Cer_NS) at 3 weeks, hexosylceramides (HexCer) at 3 and 7 weeks, sulfatides (SHexCer) at 3 weeks, and sphingomyelin (SM) at 3 and 7 weeks (**Figure 4A**). In *Npc1^−/−^* cortex Cer_NS, HexCer, and SHexCer exhibit more consistently decreased levels across all the time points, while hydroxy ceramides and gangliosides are consistently increased (**Figure 4B**). These observations point out that the cerebellum sphingolipid profile shows a disease severity-dependent trend, while most of the cortex sphingolipids seem to be consistently elevated throughout disease progression. Moreover, in the cortex, hydroxy ceramides exhibit an opposite trend compared to non-hydroxy ceramides, while in the cerebellum, hydroxy ceramides show more pronounced accumulation than their non-hydroxy counterparts. This observation is further discussed below detailing myelin changes.

**Figure 4.**
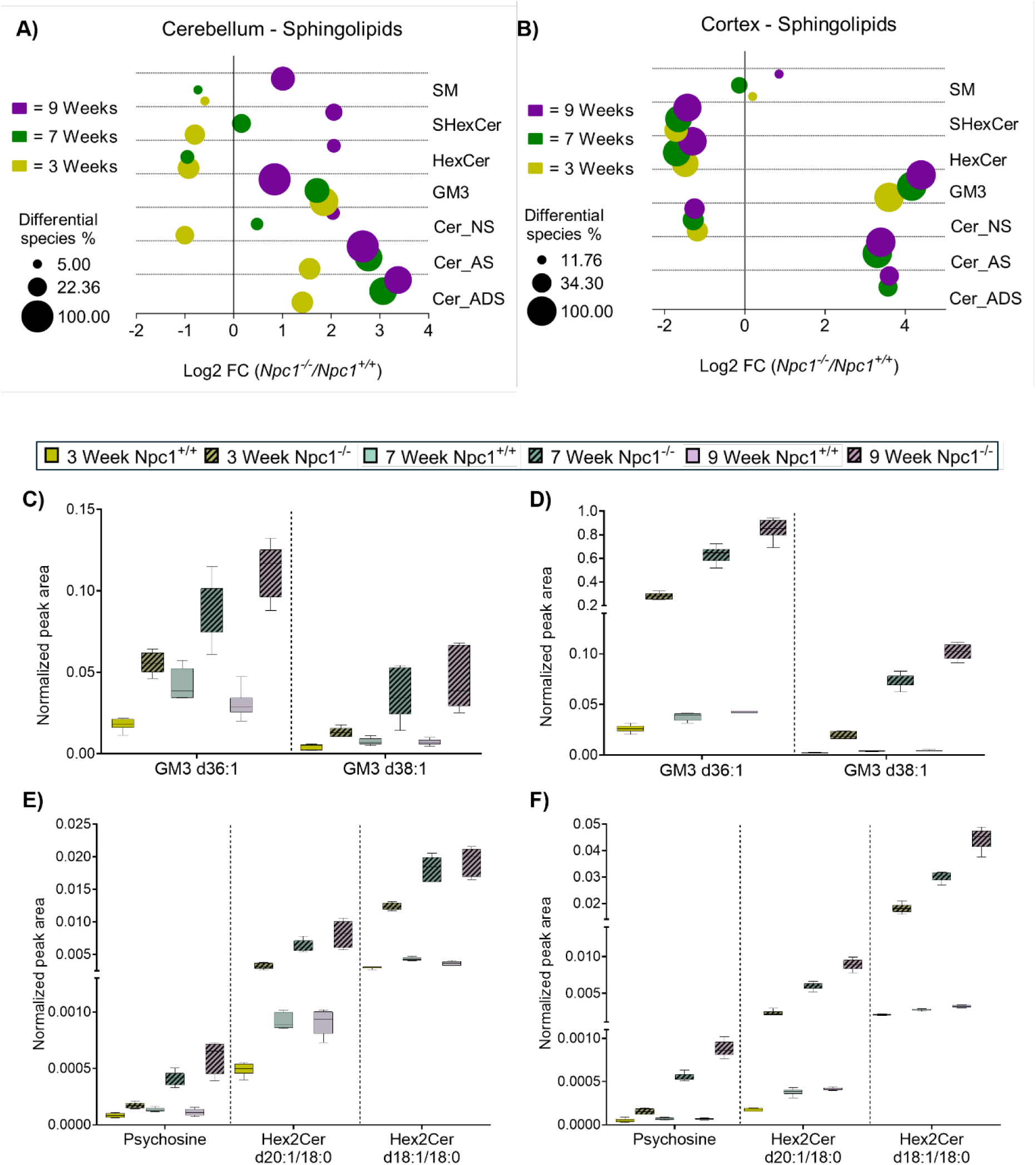
Sphingolipids show significant changes in *Npc1^−/−^* tissues. A) cerebellum and B) cortex differential sphingolipids at each time point. Summed, normalized abundances of the differential lipids were used to calculate the Log2 Fold change (FC) and used to create the bubble plots. Bubble size is scaled to represent the percentage of the significantly different lipid species compared to number of total identified lipid species in each lipid class. An Unpaired t-test was conducted between *Npc1^−/−^* and *Npc1^+/+^* for each time point and each tissue to calculate the p-values and the lipids with p-value < 0.05 and FC > 1.5 were considered significant. All the lipids included in the calculations are significantly different between the 2 groups. Ganglioside GM3 species showing severe accumulation in the *Npc1^−/−^* C) cerebellum D) cortex tissues. Psychosine and dihexosylceramide (Hex2Cer) species that show significant accumulation in the *Npc1^−/−^* E) Cerebellum F) Cortex tissues. Peak areas normalized to the internal standards were used to create the box and whisker plots.

The gangliosides are one of the most well-known glycosphingolipids that accumulate in NPC1 disease^18,34–36^ and it has also been proposed that the cholesterol accumulation in NPC1-deficient neurons is ganglioside-dependent^34^. Interestingly, while the cerebellum is the most severely affected brain region in NPC1 disease,^37^ ganglioside GM3 displays more pronounced accumulation in the cortex of *Npc1^−/−^* mice (**Figure 4A-4D)**. We report that the GM3 ganglioside accumulating in *Npc1^−/−^* mouse brain is incorporated with sphingosine backbones 18:1 and 20:1, and contains stearic acid (a long-chain saturated fatty acid) as the fatty acyl component (**Figure 4C, 4D**). It is also evident from our data that the level of accumulated GM3 in both brain regions increases significantly with disease progression (**Figure 4C-4D**). Another glycosphingolipid and a potential GM3 precursor, dihexosylceramide (Hex2Cer), also shows significant accumulation in both brain regions (**Figure 4E, 4F**). Interestingly, following the pattern seen with GM3, the Hex2Cer species also show greater accumulation in the cortex compared to the cerebellum (**Figure 4E, 4F**). In addition, the sphingosine and fatty acid composition of the altered Hex2Cer species matches that of the elevated GM3 species. This implies that the accumulating Hex2Cer species are the precursor molecules driving elevated GM3 species. Moreover, in line with previous reports^38^, we also report significant accumulation of the free sphingolipid backbone molecules, sphingosine and dihydrosphingosine (also known as sphinganine) in both brain regions **(Figure S8)**.

Expanding the knowledge of sphingolipid alteration in the NPC1 disease, we reveal that galactosylsphingosine-also known as psychosine, a single-chain glycosphingolipid with a high cytotoxicity-is significantly elevated in both cerebellum and the cortex regions of the *Npc1^−/−^* mice brain (**Figure 4E, 4F**). Psychosine accumulation becomes significant as early as the 3-week time point and appears to increase progressively with disease progression. Psychosine is a highly cytotoxic lipid even at lower concentrations and is capable of inducing cell death, in cells including oligodendrocytes^39^. Yenisleidy et al. have demonstrated that psychosine can disrupt the physical and electrostatic properties of lipid membranes, including the myelin membrane^40^. In addition, the accumulation of psychosine has been proposed to be the pathogenic mechanism as a result of dysfunctional galactosylceramidase (GALC) in Globoid cell leukodystrophy (GLD), also known as Krabbe disease^41,42^. Psychosine is synthesized by the diacylation of galactosylceramide (GalCer) by acid ceramidase and is readily degraded by GALC. In Krabbe disease, the loss of GALC activity is known to result in psychosine accumulation^43^. Interestingly, Oliver et al. have reported that the NPC1-depleted cells exhibit reduced GALC activity compared to the control cells^44^. We hypothesize that the elevation of psychosine levels in *Npc1^−/−^* mice brain observed in this study is a result of reduced GALC activity, and the cytotoxicity could be contributing to the neuronal loss observed in NPC1 disease.

### Myelination defect in *Npc1^−/−^* mouse brain

The NPC1 and NPC2 proteins are lysosomal cholesterol transporters that facilitate cholesterol egress through the endo/lysosomal compartments ^45,46^. Loss of function of these proteins leads to the accumulation of unesterified cholesterol (free cholesterol) in the lysosome^47^. Despite the cholesterol accumulation phenotype within lysosomes in NPC1 disease, our data suggest that unesterified cholesterol levels in the *Npc1^−/−^* mice brain is significantly decreased and accumulations are location specific within cellular compartments (**Figure 5**). In the cerebellum, cholesterol levels are significantly decreased at the 3- and 7-week time points and similar to wild-type levels at 9 weeks. In the cortex, significant reductions are observed at 7 and 9 weeks (**Figure 5**). We hypothesized that this observation is due to the dysmyelination (improper myelination) phenotype in the NPC1 disease^7,8,48^. In contrast to other biological membranes, where the lipid to protein ratio is close to one, the myelin membrane contains a higher proportion of lipids (70% - 85%)^49^. While no myelin-specific lipids have been reported to date, cholesterol accounts for the majority of lipid content in the myelin (more than 45% in the central nervous system)^49,50^. The reduction of myelin mass in the brain is likely to have a substantial impact on cholesterol levels as well. To confirm this, we isolated myelin from *Npc1^−/−^* and *Npc1^+/+^* mice brains at 5 and 9 weeks using a previously published density-based separation method^51^. The dry weight measurements of the isolated myelin confirm our hypothesis, showing a significant decrease in myelin mass in both cerebellum and cortex brain regions (**Figure 6A, 6B**). Moreover, the myelin loss in the cortex region seems to be more pronounced than in the cerebellum.

**Figure 5.**
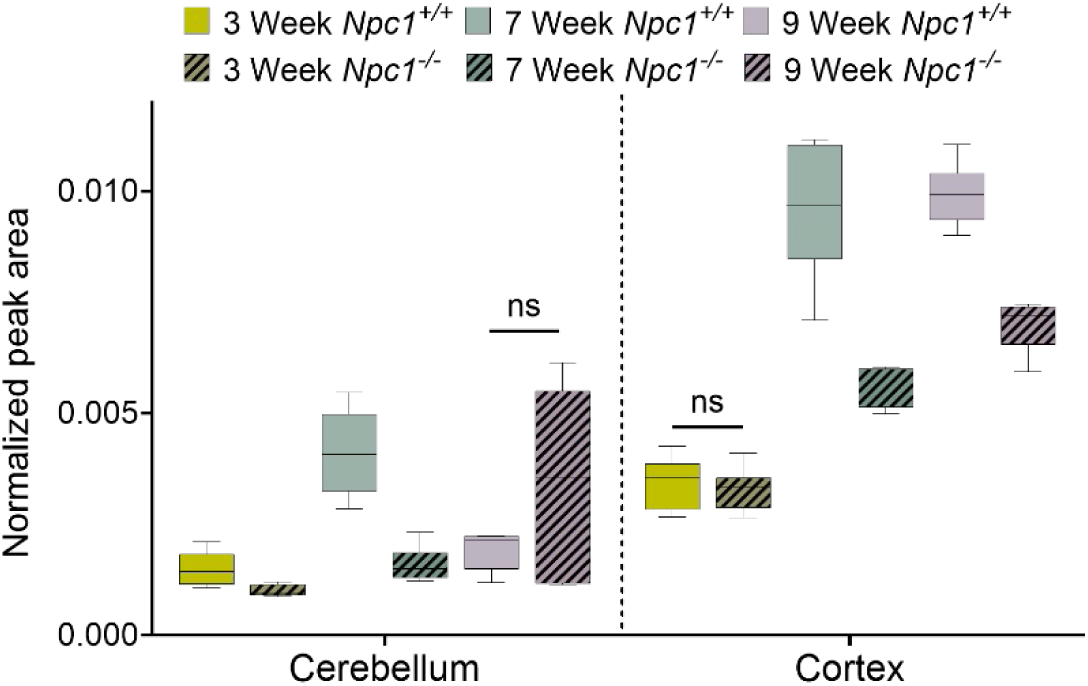
Decrease of cholesterol levels in the *Npc1^−/−^* cerebellum and cortex tissues. Peak areas normalized to the internal standards were used to create the box and whisker plots. An unpaired t-test was conducted to determine the p-value. p-value > 0.05 is considered as nonsignificant (ns)

**Figure 6.**
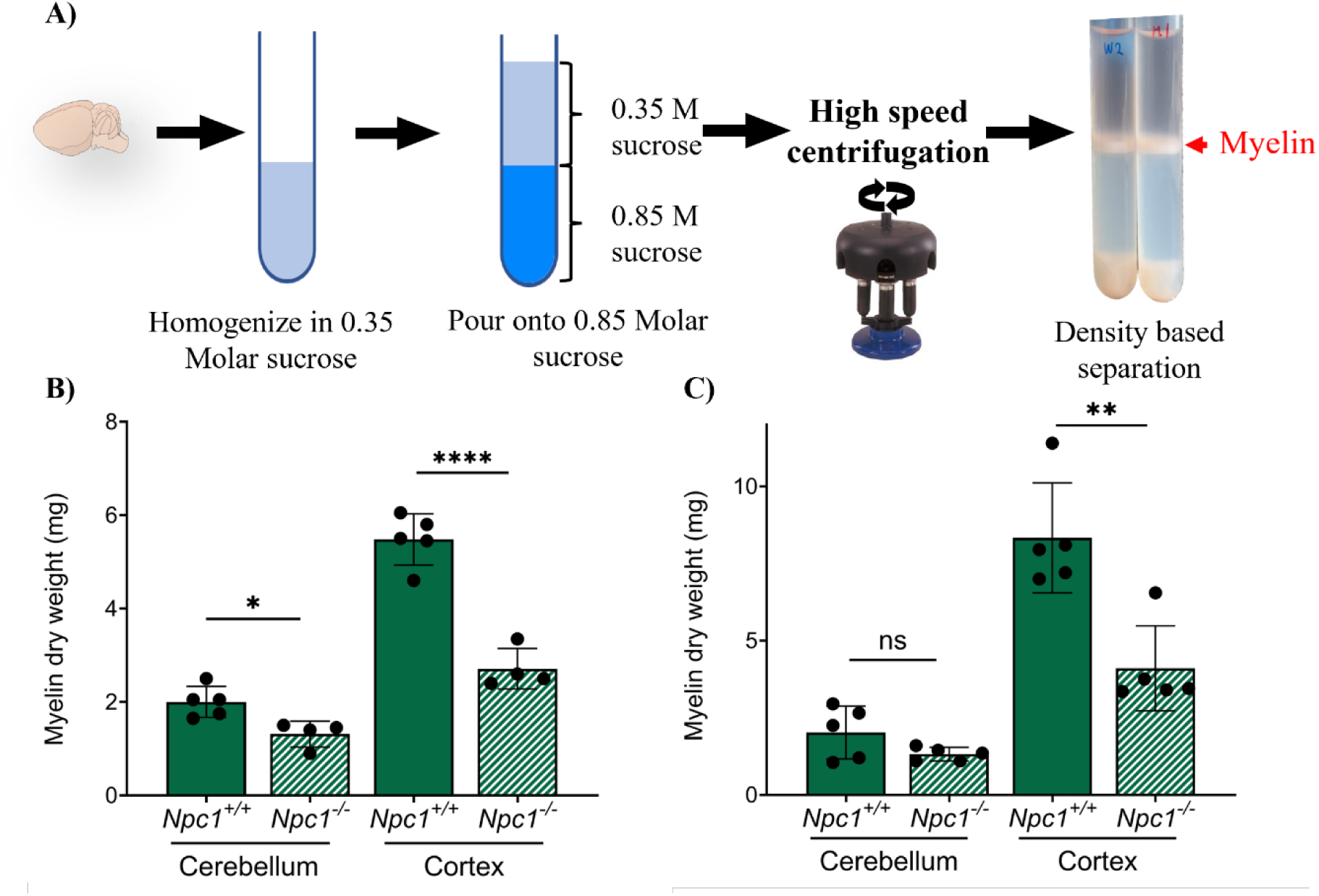
Density-based myelin isolation from mice brain cerebellum and cortex A) Workflow. Isolated myelin from each genotype at age B) 5 weeks and C) 9 weeks shows decreased myelin in the *Npc1^−/−^*. Severe dysmyelination is observed in the cortex compared to the cerebellum. Unpaired t-test was performed to determine the p-value. (p-value < 0.001, ****, p-value < 0.01, **, p-value < 0.05, *)

### Alterations in myelin lipidome of the *Npc1^−/−^* mouse cerebellum and cortex

Consistent with the reduction in cholesterol, additional predominant myelin-associated lipid species—including sulfatide (SHexCer), hexosylceramide (HexCer), phosphatidylcholine (PC), phosphatidylethanolamine (PE), ether-linked PE (PE O-), and non-hydroxy ceramide (Cer_NS)—also exhibit significant decreases. These lipid species exhibit consistent decreases across all three time points in the whole cortex, whereas in the whole cerebellum, reduced levels of these lipids are primarily observed at the earlier stages of disease progression (**Figure 2A, 2B, 4A, 4B**). Furthermore, distinct accumulation of hydroxy ceremide species, which are essential in maintaining the structural integrity and stability of myelin^52^, is observed in both brain regions (**Figure 4A, 4B**). Given the above findings and the essential role of lipids in maintaining myelin integrity and function, we systematically examined the alterations in brain myelin lipid composition resulting from NPC1 deficiency. Our results provide molecular details of dysmyelination observed in NPC1 disease.

As the first comprehensive analysis of the myelin lipidome in NPC1 disease, our study identifies distinct lipid alterations within cerebellar and cortical myelin. In general, cortical myelin exhibited a higher degree of lipid perturbations across all time points, with the severity of myelin lipid dysregulation increasing as the disease progressed (**Figure 7A, 7B, 8A**). Myelin lipidomic results and the isolated myelin content measurements imply that the myelin in the cortex region is more severely affected compared to cerebellar myelin in the NPC1 disease. In cortical myelin, a general trend of increased lipid species is seen, where some lipid classes such as phosphatidylcholine (PC), phosphatidylethanolamine (PE) and phosphatidylserine (PS) exhibit mixed trends (**Figure 7A, 7B)**. However, lipids including acylcarnitines (Acar), sulfatide (SHexCer), and ether PE (PE O-), show a consistent decrease at both time points (**Figure 7A, 7B)**. The glycosphingolipid, sulfatide is one of the major lipids found in myelin, comprising 4% of total myelin lipids^53^. Beyond their role as structural lipids, sulfatides are also key molecules in maintaining ion channels on myelinated axons^54^. More importantly, sulfatides are implicated in oligodendrocyte differentiation, a process known to be impaired in NPC1 disease^7,55^. Further studies are required to determine whether sulfatide loss in NPC1 disease contributes to the myelin ion channel activity or the dysmyelination phenotype observed in the central nervous system. Manipulation of the sulfatide level could be a potential approach to rescue the dysmyelination in NPC1 disease.

**Figure 7.**
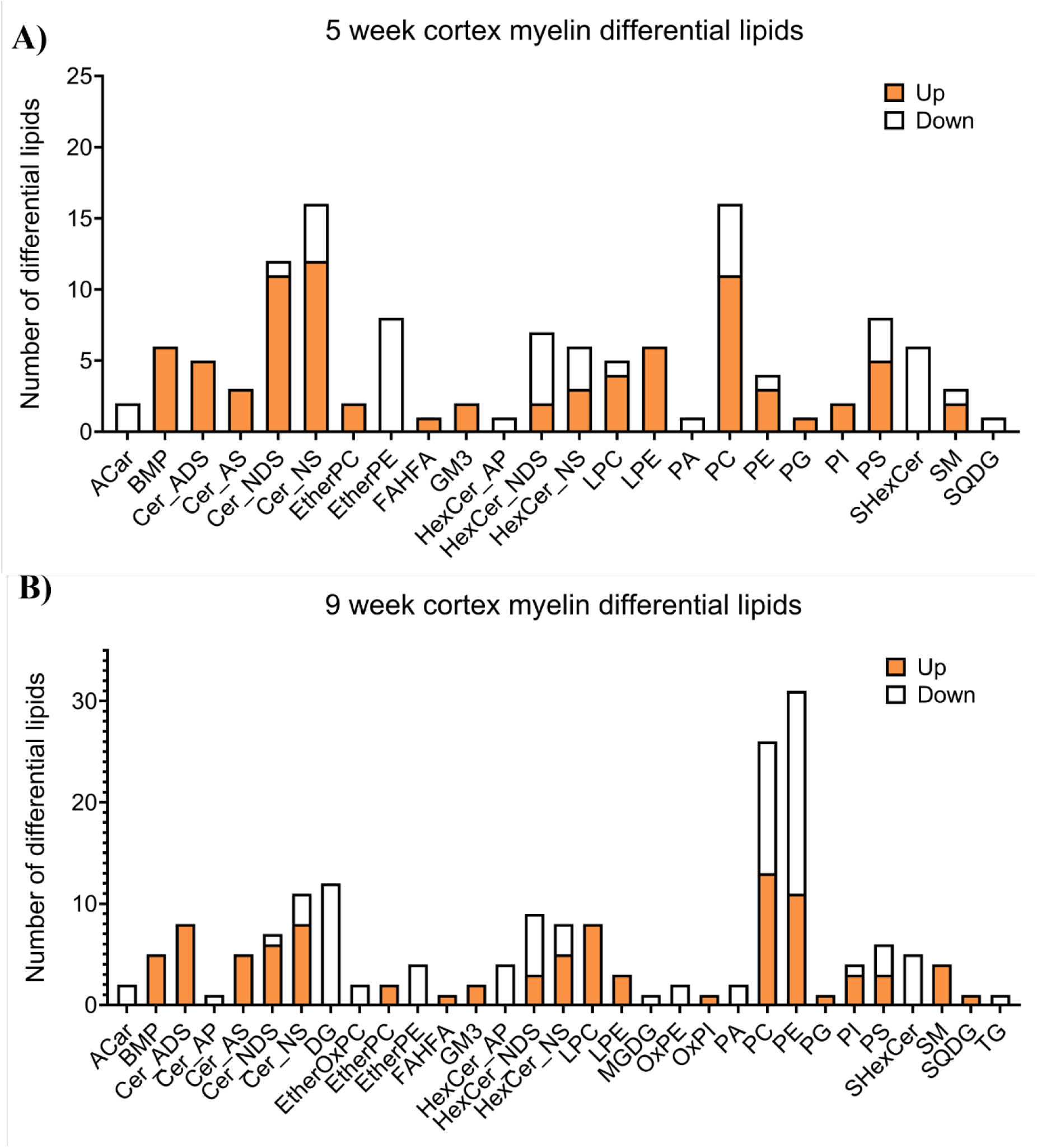
Differential lipids in the myelin isolated from *Npc1^−/−^* mice cortex region at A) 5 weeks, and B) 9 weeks time points. Number of differential lipid species per lipid class is plotted in the y-axis. Number of accumulated lipid species is shown in orange color and the depleted lipids are represented by white color. Unpaired t-tests were conducted between *Npc1^−/−^* and *Npc1^+/+^* for each time point and each tissue to calculate the p-values and the lipid changes with a p-value < 0.05 and FC > 1.5 were considered significant.

In reference to the whole-tissue lipidomic data presented above, myelin lipidomics revealed distinct lipid alterations, particularly at early time points in the cerebellum. In the cerebellar whole-tissue lipidomic data, lipid classes including sphingomyelin (SM), phosphatidylserine (PS), phosphatidylethanolamine (PE), phosphatidylcholine (PC), and hexosylceramide (HexCer_NS) showed substantial decreases at the 3-week and 7-week time points. However, the 5-week myelin lipidomic data revealed that these lipid classes were either significantly increased or do not show a significant change in cerebellar myelin (**Figure 8A**). A similar observation was also made in cortical myelin, where lipid classes including lysophosphatidylcholine (LPC), PS, phosphatidylinositol (PI), HexCer_NS, and non-hydroxy ceramide (Cer_NS) showed comparable changes (**Figure 8B**). This observation supports our hypothesis that the lipid reduction observed in whole-tissue samples is likely due to myelin loss. It highlights the importance of investigating the myelin lipidome and regional changes to determine whether the disease condition truly impacts lipid composition.

**Figure 8.**
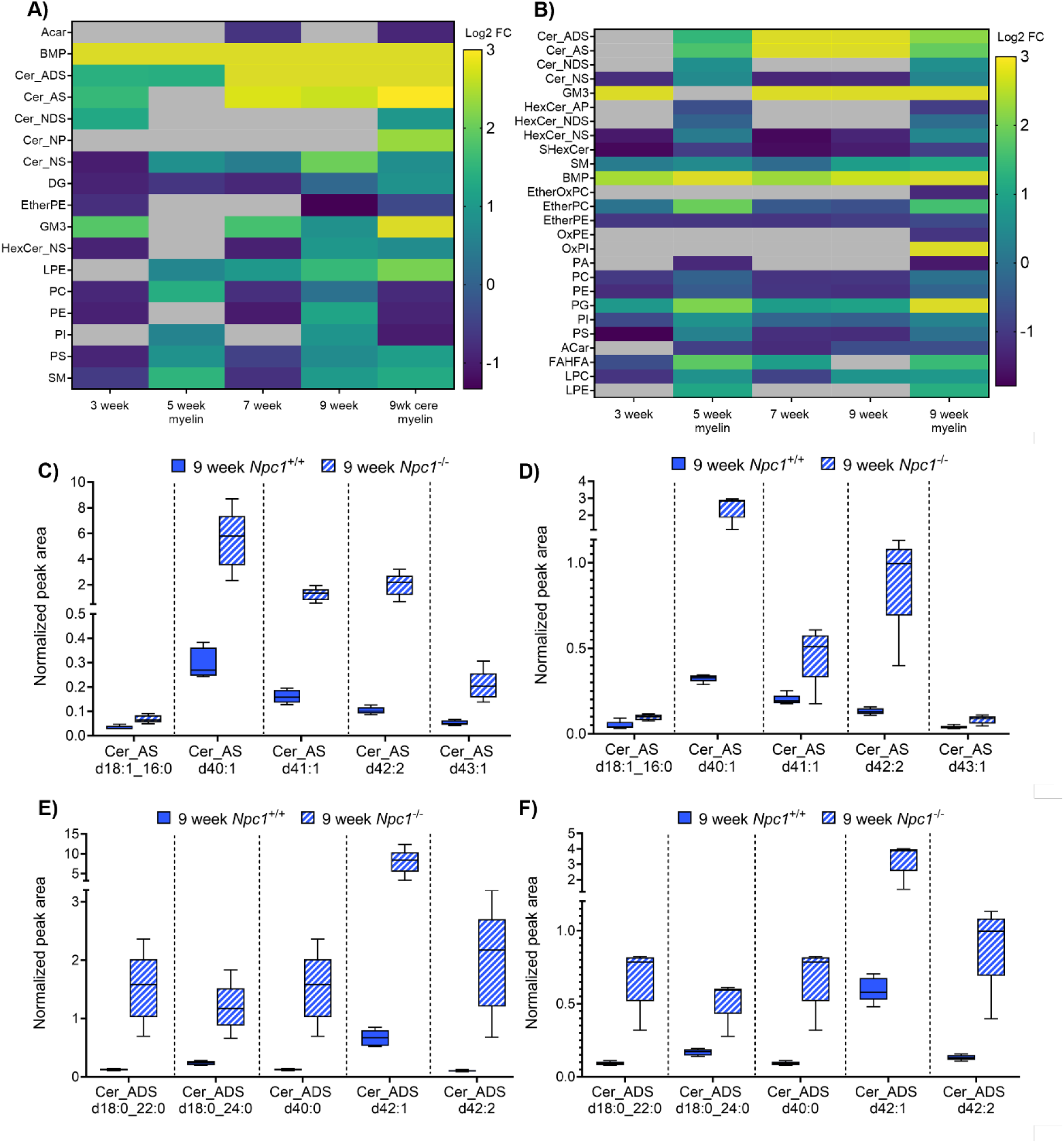
Myelin lipid alteration of each lipid class. Heatmap showing the differential lipids in the isolated myelin along with the regulation of these lipids in whole tissue A) Cerebellum. B) Cortex. An Unpaired t-test was performed to determine the p-value, and lipid changes with p-value < 0.05 and FC > 1.5 are considered significant. Lipids that did not show significant changes are greyed in the heatmap. Log2 FC of the significant lipids are used to assign a color from the color scale. Bright yellow is used to show the lipids with a Log2 FC > 3. Hydroxy ceramide accumulation in the 9-week cerebellum and cortex myelin, peak area normalized to internal standards, is used to create graphs for the alpha-hydroxy fatty acid ceramides (Cer_AS) in C) cerebellum myelin D) cortex myelin and alpha-hydroxy fatty acid sphingonine ceramide (Cer_ADS) in E) cerebellum myelin F) cortex myelin.

Lastly, among the most prominent lipid changes in the myelin lipidome, alterations in hydroxy ceramides represent a particularly significant finding. While multiple hydroxy ceramide lipid species exhibited significant accumulation at both time points, the 9-week myelin data indicate a more pronounced accumulation (**Figure 8A, 8B**). At the 9-week time point, alpha-hydroxy fatty acid ceramides (Cer_AS) and alpha-hydroxy fatty acid sphingonine ceramide (Cer_ADS), also known as 2-hydroxy ceramide, exhibit significant accumulation in both cerebellar and cortical myelin (**Figure 8C-8F**). Some of these altered hydroxy ceramide lipid species show a 10 to20-fold increase compared to the wild-type myelin. Hydroxy sphingolipids are considered to play an essential role in myelin structure and function^56^. This is evident in fatty acid 2-hydroxylase (FA2H)-deficient models, where FA2H is responsible for the synthesis of 2-hydroxylated sphingolipids^56,57^. Studies have shown that the presence of the hydroxyl group near the polar head group in these lipids facilitates a hydrogen bonding network within neighboring lipids to improve membrane rigidity^58,59^. Given the importance and unique properties of hydroxy sphingolipid in the myelin membrane, the severe accumulation of these lipids in NPC1 myelin could directly affect the physical properties of myelin. Further studies are needed to explore the impact of hydroxy ceramide accumulation on myelin properties and neuronal function in NPC1 disease.

## Discussion

The *Npc1^−/−^* mice model used in the current study recapitulates many of the neuropathological features of the human disorder, including cerebellar Purkinje cell loss and neuronal lipid storage, ^3,16^. Sphingolipids are well known to be altered in NPC1 disease and have been characterized in the whole brain of the *Npc1^−/−^*mouse model^18^. In this study, we employed an LC-MS-based discovery lipidomic approach to investigate the lipidome of the cerebellum and cortex of the *Npc1^−/−^* mouse model in an unbiased manner. Based on our findings in the whole tissue, we broadened our study to investigate the myelin lipidome in *Npc1^−/−^* mouse model and reveal significant lipid perturbations in both cerebellar and cortical myelin.

We found significant changes in glycerophospholipids in the *Npc1^−/−^*mouse. Among the differential glycerophospholipids, the bis(monoacylglycero)phosphate (BMP) lipid accumulation is particularly important due to its localization and critical functions within the lysosome. The data also indicate that the elevated BMP lipids are enriched with docosahexaenoic acid (DHA). This elevated DHA incorporation to BMP could be reducing the accessible free DHA levels in the lysosome, which is critical in lysosomal autophagy^60,61^. Importantly, DHA and its metabolites in the brain play a crucial role in brain growth, development, and the modulation of glial cells, and are a major lipid enriched in neuronal membranes ^62^. Furthermore, intermediate lipids in the BMP biosynthesis pathway,lysophosphatidylglycerol (LPG) and hemi-BMP (HBMP) species, with fatty acid compositions matching those of the accumulating BMP lipids, are also elevated in both brain regions as early as the 3-week time point before disease symptom onset. The accumulation of BMP synthesis intermediates suggests that the biosynthetic pathway is impaired following the loss of NPC1 activity in the mouse brain. This could be due to increased lysosomal pH in NPC1 disease, which may reduce the efficiency of enzymes like lysosomal PLA2 and CLN5 that are involved in BMP synthesis^30,63^. This finding is particularly crucial in NPC1 disease due to the involvement of BMP lipids in cholesterol trafficking and growing interest in understanding BMP biology^22,33,64^. Studies have demonstrated that the supplementation of BMP and hydrolysable phosphatidylglycerol (PG) lipids with mono and polyunsaturated long-chain fatty acids resulted in cholesterol clearance from the NPC1-deficient cells^22,65^. However, the mechanism by which BMPs facilitate cholesterol clearance from NPC1-deficient cells is not clear. The increased levels of endogenous BMPs observed in the mouse could be due to dysfunction of lysosomal phospholipase PLA2G15^64^, or there may be a feedback mechanism to stimulate BMP production in an effort to clear cholesterol accumulation. However, the elevated endogenous BMP levels in NPC1-deficient cells may be insufficient to facilitate cholesterol clearance, or they may not be functionally accessible to participate in BMP-induced cholesterol clearance. Additional work is needed to define if our observation is broadly applicable to other tissues or just the brain in NPC1.

Consistent with former studies in this mouse model, we report multiple sphingolipid species are elevated in both cerebellum and cortex regions^18^. Expanding the current knowledge in sphingolipid defects in NPC1 disease, our data revealed the progressive accumulation of the galactosylsphingosine, also known as psychosine. The psychosine accumulation is a hallmark in Globoid cell leukodystrophy (GLD), also known as Krabbe disease, in which severe demyelination is reported^66^. This cytotoxic lipid is known to cause an inflammatory response and cell death in the CNS, especially in astrocytes and myelin-forming oligodendrocytes^39,67^. A mechanistic study on psychosine toxicity in a mouse-derived oligodendrocyte progenitor cell line revealed activation of caspase-9 and downstream caspases, suggesting that psychosine induces cell death via apoptosis^68^. Interestingly, in NPC, while differing reports describe cell death pathways, the neuronal cell death has been shown to be caspase-dependent, suggesting the potential contribution of psychosine accumulation to neuronal cell death in NPC^69^. The psychosine toxicity in oligodendrocytes could be contributing to the dysmyelination phenotype seen in NPC1 disease and should be further explored. Furthermore, targeting the psychosine production represents a potential therapeutic strategy in NPC1 disease, aiming to mitigate psychosine-induced cytotoxicity and neuroinflammation.

Analysis of the lipidome of isolated myelin from the cerebellum and cortex of NPC1 mice revealed a significant reduction in myelin dry mass in both brain regions of *Npc1⁻^/^⁻* mice and highlights the more pronounced reduction of myelin in the cortex region. This observation is consistent with the reduced myelin basic protein (MBP) expression observed in the cortex reported in a previous study^7^. The dysmyelination phenotype in NPC1 has been investigated in multiple studies, and inhibition of oligodendrocyte maturation has been observed upon the loss of NPC1 protein function^70^. Isolated myelin from cerebellum and cortex regions was successfully used to conduct LC-MS/MS lipidomic analysis and revealed multiple lipid alterations in both brain regions; however, a significantly greater number of lipids and higher fold changes were observed in the cortex region. The observed lipid perturbations in this study, such as decreased sulfatide and cholesterol levels, could play a significant role in the oligodendrocyte differentiation defect reported in NPC1 disease^71^. Providing evidence that sulfatide lipids are also important in myelin maintenance and survival, studies have shown that mild progressive demyelination is observed in a ceramide sulfotransferase knockout mouse model^72,73^. In addition, myelin structural lipid alterations, such as 2-hydroxy sphingolipids, could impact the ordered myelin structure, resulting in poor myelin rigidity and compactness. Overall, our study points out the importance of investigating the myelin lipidome in demyelination and dysmyelination disorders. However, future studies will need to investigate if manipulation of lipid synthesis and metabolism to normalize these lipid changes could rescue the dysmyelination in NPC1.

## Conclusion

The brain lipidome study carried out in NPC1 null mouse brain has provided a thorough analysis of the impact of the disease on the brain lipidome. Specifically, revealing novel lipid changes in BMP lipid synthesis intermediates, LPG and HBMP, and cytotoxic psychosine lipid in both brain regions. We provide evidence of the unique alteration trend in the cerebellar lipidome compared to that of the cortex during NPC1 disease progression. The reduced myelination in *Npc1⁻^/^⁻* mice is shown by isolating myelin from the brain, and further lipidomic analysis reveals numerous myelin lipid defects, including sulfatide and 2-hydroxy sphingolipids. These findings enhance our understanding of the lipidome in NPC1 disease and will be instrumental in guiding future research efforts. Furture in vitro and in vivo studies will need to be carried out to investigate the potential of currently available and potential treatment strategies to normalize the reported lipid perturbations.

## Materials and methods

### Materials

Methanol (LC-MS grade), acetonitrile (LC-MS grade), chloroform (LC-MS grade), ammonium acetate (reagent grade), phenylmethylsulfonyl fluoride (PMSF), sodium fluoride (NaF), sucrose (molecular biology grade), and sodium orthovanadate (Na3VO3) were obtained from Sigma-Aldrich, St. Louis, MO. Isopropanol (LC-MS grade), Pierce BCA Protein Assay (P.N. 23227) and 10X Phosphate Buffered Saline (PBS) were obtained from Thermo Fisher Scientific, Waltham, MA. the BeatBoxTM tissue homogenizer was obtained from PreOmics, Munich, Germany. EquiSPLASH™ LIPIDOMIX™ Quantitative Mass Spec Internal Standard (P.N. 330731) was obtained from Avanti Research, Birmingham, AL. Pestle homogenizer (Homogenizer motor/control dual shaft) was obtained from Glas-Col, Terre Haute, IN. Probe sonication was obtained from Qsonica sonicator, Newtown, CT. The SW 41 Ti swinging rotor was obtained from Beckman Coulter, Brea, CA. All reagents were used as supplied unless otherwise noted.

### Animal Model

All experiments were conducted in accordance with the University of Illinois Chicago IACUC-approved protocol (ACC Protocol Number: 24-102). The Balb/c *Npc1* heterozygote (*Npc1^+/−^*) mice were obtained from The Jackson Laboratory (RRID:IMSR JAX:003092), and a breeding colony was maintained in our laboratory. Genotyping was performed using PCR as previously reported ^74^. At 3, 7 and 9-weeks of age, control (*Npc1^+/+^*) and null *(Npc1^−/−^*) mice were euthanized via CO2 asphyxiation followed by decapitation. Whole brain tissue was dissected, immediately frozen in dry ice to maintain spatial integrity and stored at −80 °C.

### Lipid extraction from mouse cerebellum and cortex for LC-MS analysis

Cerebellar and cortex tissues were homogenized in 1X PBS buffer with phosphatase inhibitors (1 mM NaF, 1 mM β-Glycerophosphate, 1 mM PMSF, 1 mM Na3VO3) using the BeatBox^TM^ tissue homogenizer. For the homogenization, tissue samples (40 – 50 mg) were placed in a 2 mL BeatBox tissue kit 24× tube (P.O. 00128), and four rounds of 5-minute standard cycles were run. Between each 5-minute cycle, samples were cooled on ice. The total protein content of the lysed samples was determined using the Pierce BCA Protein Assay. To an equivalent of 70 µg of protein from each sample, 1X PBS was added to the final volume of 40 µL followed by 5 µL of the EquiSPLASH™ LIPIDOMIX™ Quantitative Mass Spec Internal Standard (P.N. 330731). To each sample, 800 µL of the ice-cold CHCl3: Methanol (2:1, v/v) was added and vortexed. Samples were then incubated on ice for 15 minutes, while vortexing every 5 minutes. After the incubation time, 160 µL of 1X PBS was added and vortexed, followed by centrifugation at 1500 rpm for 5 minutes at room temperature to induce phase separation. The upper organic phase was transferred into a fresh tube and dried under vacuum. Dried samples were resuspended in 70 µL of methanol/CHCl3 (3:1, v/v) and centrifuged at 4 °C, 14,000 rpm for 10 minutes before transferring into LC-MS vials. A pool lipid extract sample was made for each cerebellum and cortex by combining 7 µL from each individual lipid extract.

### Myelin isolation and lipid extraction for LC-MS analysis

Myelin was isolated as previously described ^50^. In brief, the tissue was homogenized in a 0.35 M sucrose solution using a pestle homogenizer. Homogenized lysate was carefully poured onto the 0.85 M sucrose layer. Samples were centrifuged at 75,000 g for 35 minutes in a SW 41 Ti swinging rotor. Myelin was collected at the interface of the two sucrose layers and further purified by an osmotic shock in water. The final purified myelin was dried under vacuum and weighed. Purified myelin 50 µg from each sample was homogenized in 1X PBS with phosphatase inhibitors by probe sonication (three rounds of 10-second pulses at 40% amplitude, 1 second on 1 second off). Following the homogenization, lipid extraction from myelin samples was conducted following the protocol described above with minor modifications: SPLASH™ LIPIDOMIX™ Quantitative Mass Spec Internal Standard 4 µL was used per sample, Final lipid extract was resuspended in 40 µL of methanol/CHCl3 (3:1, v/v).

### LC-MS analysis

An Agilent 1260 HPLC system outfitted with a Poroshell 120 EC-C18 column (2.1 × 100 mm, 2.7 μm, P.N 695775-902, Agilent Technologies, Santa Clara, CA) was used for the lipid separation. Mobile phase solvents consisting of (A) water: methanol (9:1, v/v) with 5 mM ammonium acetate and (B) isopropanol: methanol: acetonitrile (5:3:2, v/v) with 5 mM ammonium acetate for chromatographic separation. Mobile phase solvent flow rate was set to 300 µL per minute while the column was maintained at 50 °C in the column compartment. The chromatographic gradient is as follows: 70% B at 0-1 min, 70% B to 86% B from 1 – 3.5 minutes, at 86% B for 3.5 - 10 minutes, 86% B to 100% B from 10 – 11 minutes, at 100% B for 11 – 17 minutes, 100% B to 70% B from 17 – 17.1 min, and the column was equilibrated at 70% B for 10 minutes before the next run. An Agilent 6550 iFunnel Q-TOF (Agilent Technologies, Santa Clara, CA) was used for the MS analysis. Agilent Jet Stream (AJS) Electrospray ionization (ESI) source parameters were as follows: gas temp (200°C), drying gas (11 L/min), nebulizer (35 psi), sheath gas temp (250°C), sheath gas flow (12 L/min), VCap (3500 V), and fragmentor (145 V). Data acquisition was performed in both positive and negative ion modes, with an m/z range of 115 – 1700 in MS-only mode. Lipid extracts resuspended in methanol/CHCl3 (3:1, v/v) were analyzed in both positive and negative ion MS only modes. For the positive ion mode, 0.7 µL of the sample was injected, while 1 µL was injected in negative ion mode. A pooled lipid extract made by combining equal amounts of biological samples was analyzed in data-dependent acquisition (DDA) iterative MS/MS mode with a fixed collision energy of 25 eV. For the positive ion mode, 1 µL and the negative ion mode, 1.3 µL were injected, and seven iterations were collected.

### Data analysis

The DDA iterative MS/MS data from the pool samples were searched using the Agilent Lipid Annotator software for lipid identification. Search parameters used for the lipid annotator search: feature filter Q-score > 30, mass threshold ≤ 10 ppm, fragmentation score ≥ 30, total score ≥ 60. A database of identified lipids was created, and Agilent Profinder (version 10.0 sp1) software was used to generate and integrate extracted ion chromatograms from each sample. The following are the parameters used for targeted feature extraction: Match tolerance for *m/z* = 10 ppm, match tolerance for retention time = 0.3 minutes, *m/z* expansion for chromatographic extraction = 20 ppm, retention time expansion for chromatographic extraction = 0.5 minutes, absolute height peak filter = 5000 counts. Peak integration for each lipid and sample was manually confirmed, and manual integration correction was applied when necessary. Lipid class-based normalization and statistical analysis were conducted using the lipidomic workflow in the Agilent MassHunter workstation Mass Profiler Professional 15.1 software. The generation of bubble plots, bar charts, and further statistical analysis was conducted using the Prism-GraphPad (https://www.graphpad.com/).

## Data availability

The mass spectrometry raw data collected in this study are deposited in the public MassIVE data repository under accession code MSV000100094.

## Source data

Source data for each figure is included with the manuscript.

## Acknowledgements

This research was funded in part by the National Science Foundation (CAREER #2143920), The Ara Parseghian Medical Research Fund at Notre Dame, the National Institutes of Health (R01NS124784).

## Author contributions

Conceptualization: S.M.C. and K.C.P. Methodology: K.C.P. Investigation: K.C.P. and S.M.C. Formal analysis: K.C.P. Visualization: K.C.P. Funding acquisition: S.M.C. Project administration: S.M.C. Supervision: S.M.C. Writing – original draft: K.C.P. Writing – review & editing: K.C.P. and S.M.C.

