## Supplemental Information for "Unveiling Lipid Dysregulation: Lipidomics of Mouse Brain and Isolated Myelin in Niemann–Pick Disease Type C1"

.

Affiliations:

1. Department of Chemistry, University of Illinois Chicago, Chicago, IL 60607, USA
2. Laboratory for Integrative Neuroscience, University of Illinois Chicago, Chicago, IL 60607, USA

Correspondence:
Stephanie M. Cologna, Ph.D.
Professor
Department of Chemistry and Laboratory for Integrative Neuroscience
University of Illinois Chicago
845 W Taylor Street
MC 111 Room 4500
Chicago, IL 60607
P: 312-413-2762
E:

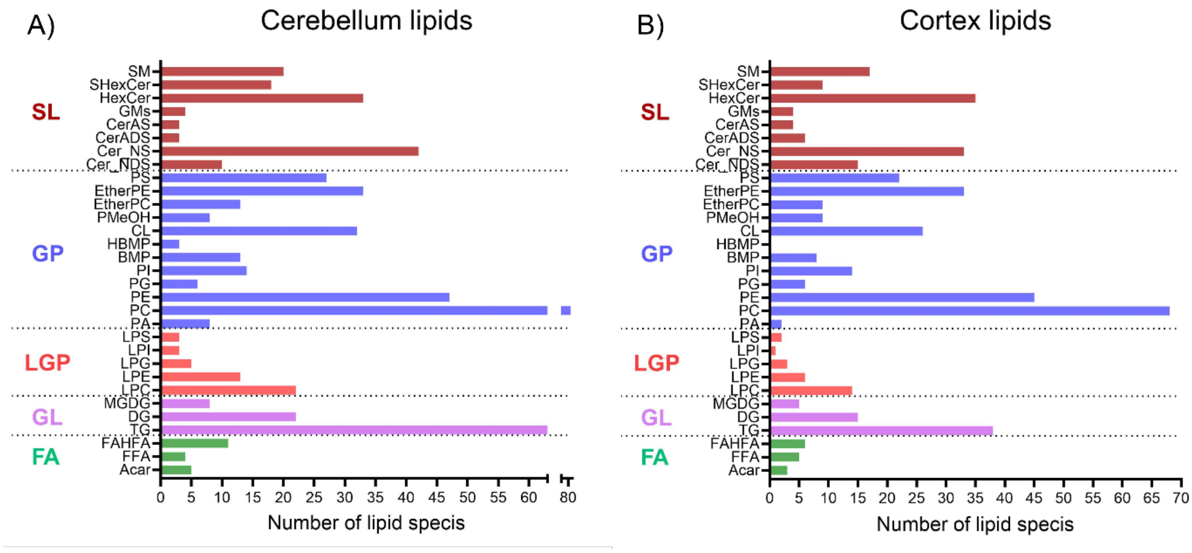


**Figure S1**. Lipid profiles of the mouse brain cerebellum and cortex. Lipids are color coded into each lipid category, and the number of species identified for each lipid class is plotted on the x-axis. Abbreviations for the lipid categories (FA = Fatty acyls, GL = Glycerolipids, LGP = Lyso-glycerolipids, GP = Glycerophospholipids, SL = Sphingolipids). Abbreviations for the lipid classes (SM = Sphingomyelin, SHexCer = Sulfatide, HexCer = Hexosyl ceramide, GM = Ganglioside, CerAS = Alpha-hydroxy fatty acid ceramides, CerADS = Alpha-hydroxy fatty acid sphingonine ceramide, Cer_NS = Non-hydroxy fatty acid ceramides, Cer_NDS = Non-hydroxy fatty acid dihydrosphingosine ceramides, PS = Phosphatidylserine, EtherPE = Ether-linked phosphatidylethanolamine, EtherPC = Ether-linked phosphatidylcholine, PMeOH = Phosphatidylmethanol, CL = Cardiolipin , HBMP = Hemibismonoacylglycerophosphate, BMP = Bis(monoacylglycerol)phosphate, PI = Phosphatidylinositol, PG = Phosphatidylglycerol, PE = Phosphatidylethanolamine , PC = Phosphatidylcholine, PA = Phosphatidic acid, LPS = Lysophosphatidylserine, LPI = Lyso-phosphatidylinositols, LPG = Lysophosphatidylglycerol, LPE = Lysophosphatidylethanolamine, LPC = Lysophosphatidylcholine, MGDG = monogalactosyldiacylglycerol, DG = Diacylglycerols, TG = Triglycerides, FAHFA = Fatty Acyl esters of Hydroxy Fatty Acids, FFA = Free fatty acids, Acar = Acylcarnitines)

A)

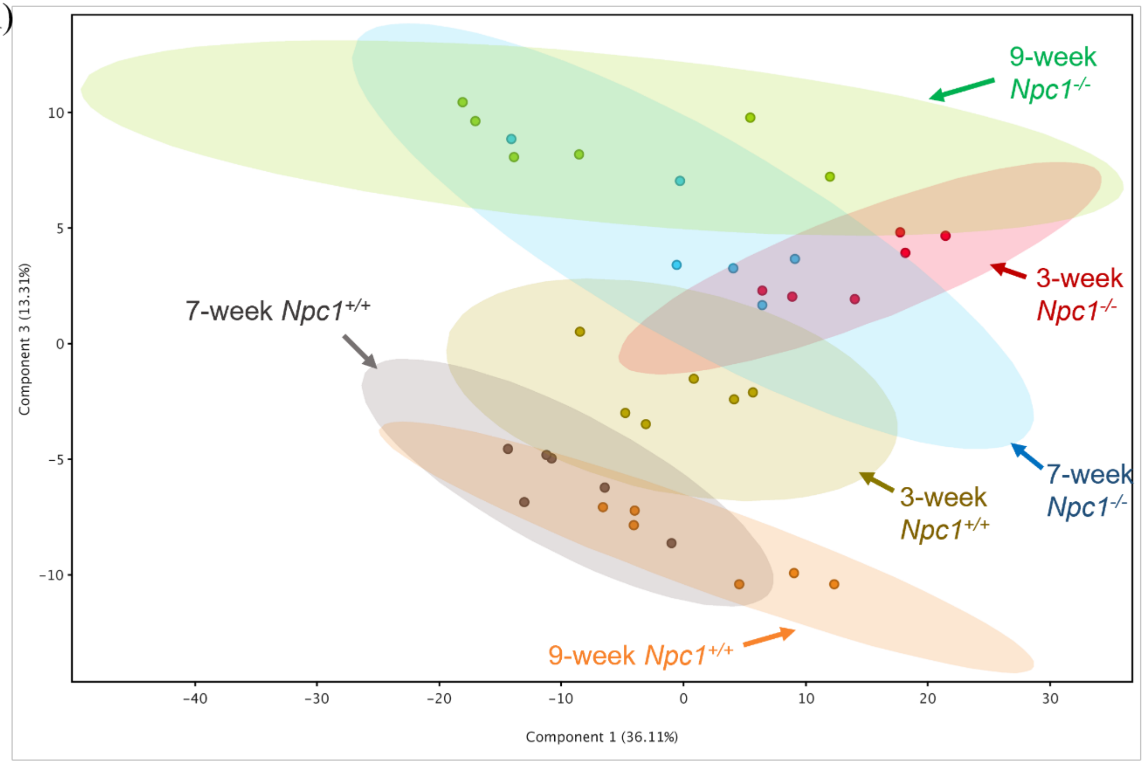


B)

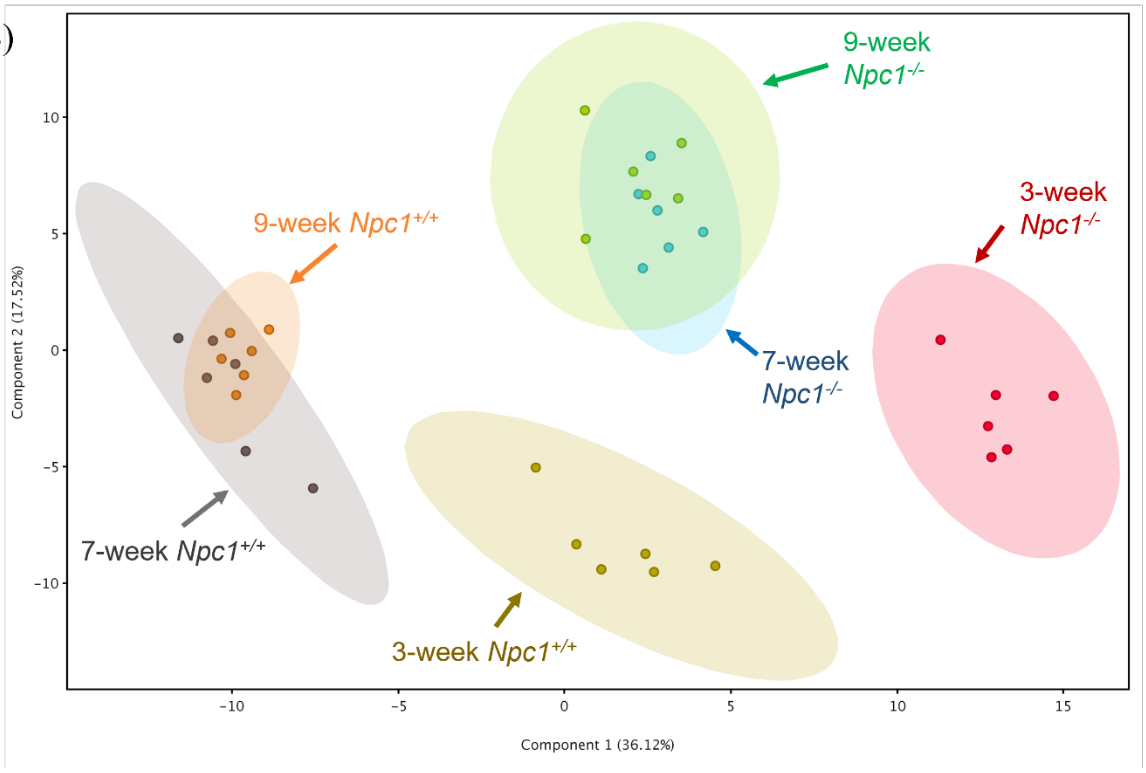


**Figure S2.** Principal component analysis (PCA) score plots showing the grouping of the biological replicates in each group and clear separation of the *Npc1*$^{+/+}$ and *Npc1*$^{-/-}$ in both A) cerebellum and B) cortex.

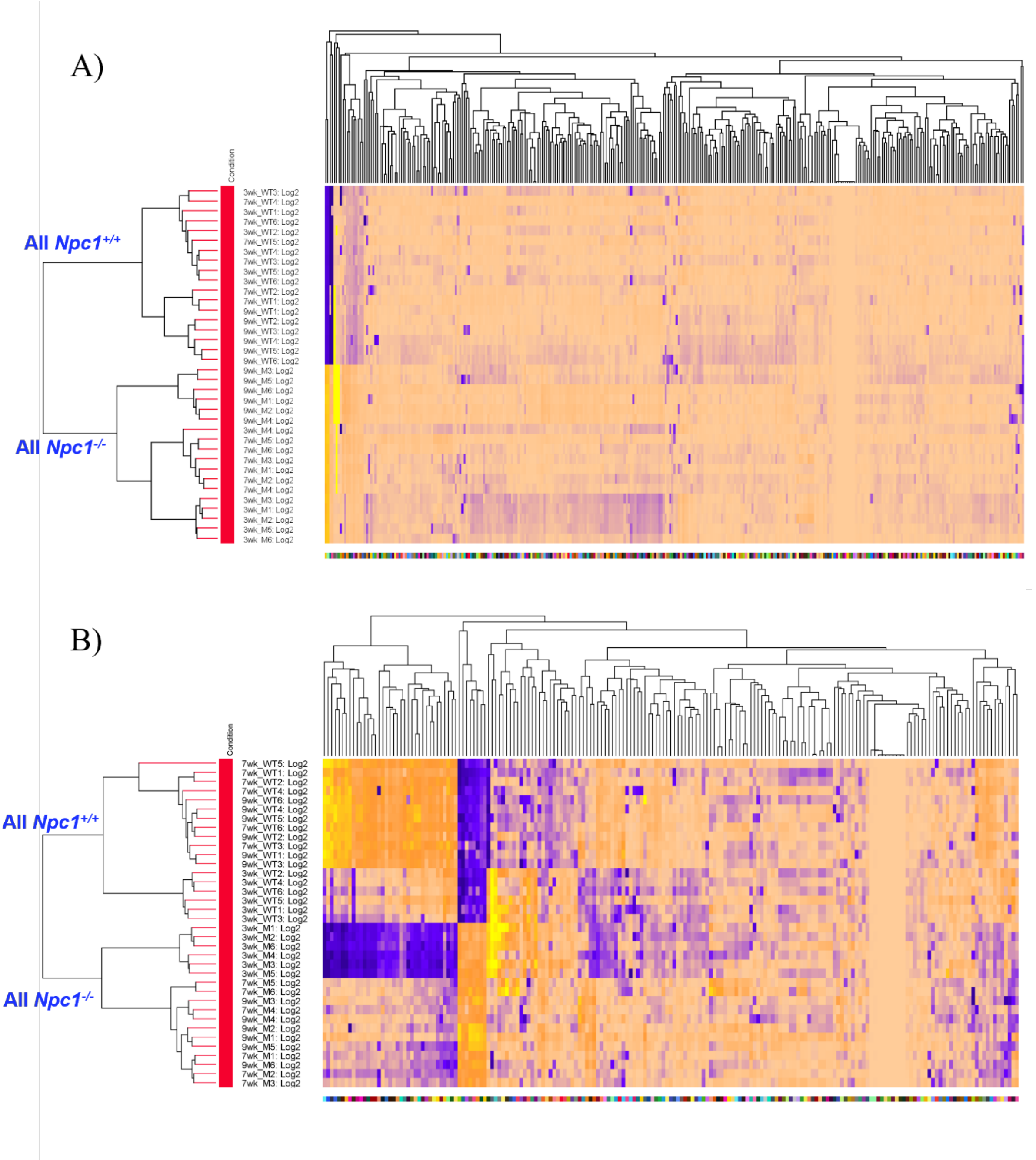


**Figure S3**. Hierarchical clustering of identified lipids by entity and by the condition shows a clear separation of the *Npc1*$^{+/+}$ and *Npc1*$^{-/-}$ in both A) cerebellum and B) cortex.

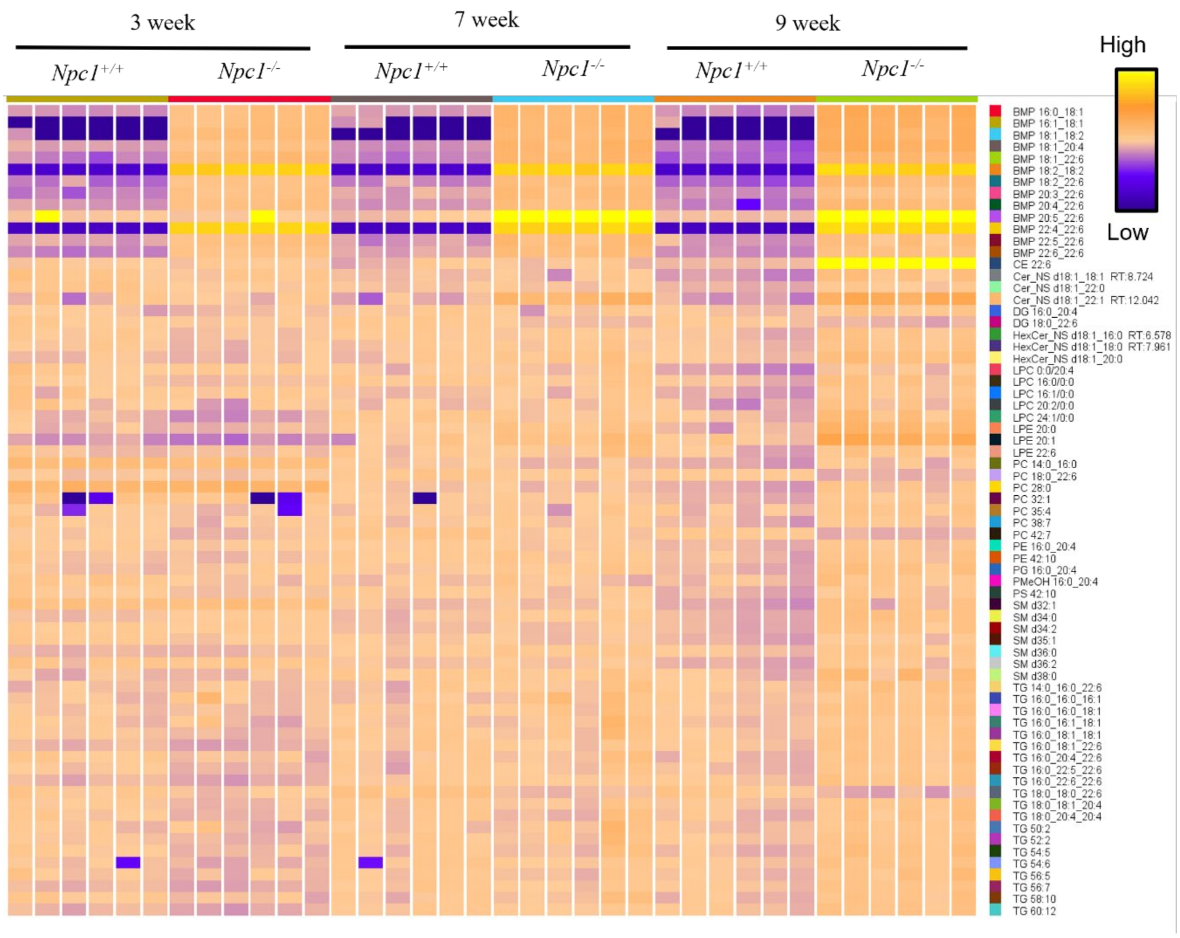


**Figure S4**. Heat map representing the lipids that are significantly different between 9-week *Npc1*$^{+/+}$ and *Npc1*$^{-/-}$ in the cerebellum. Unpaired t-tests were conducted between *Npc1*$^{-/-}$ and *Npc1*$^{+/+}$ for 9-week cerebellum positive ion mode data, and p-value < 0.05 and FC > 1.5 were considered significant. The color scale represents the normalized and baselined Log2 peak area of each lipid

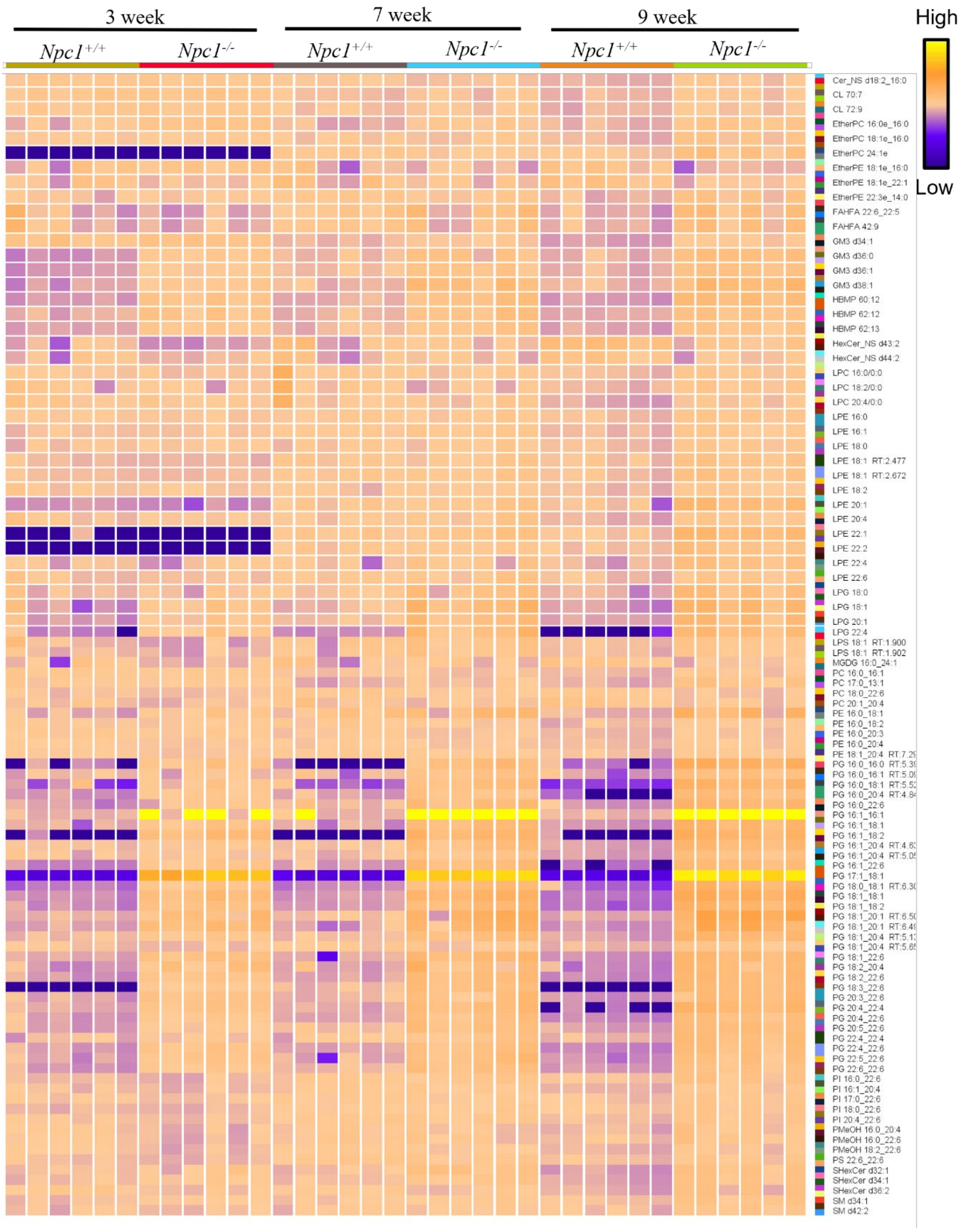


**Figure S5**. Heat map representing the lipids that are significantly different between 9-week *Npc1*$^{+/+}$ and *Npc1*$^{-/-}$ in the cerebellum. Unpaired t-tests were conducted between *Npc1*$^{-/-}$ and *Npc1*$^{+/+}$ for 9-week cerebellum negative ion mode data, and p-value < 0.05 and FC > 1.5 were considered significant. The color scale represents the normalized and baselined Log2 peak area of each lipid.

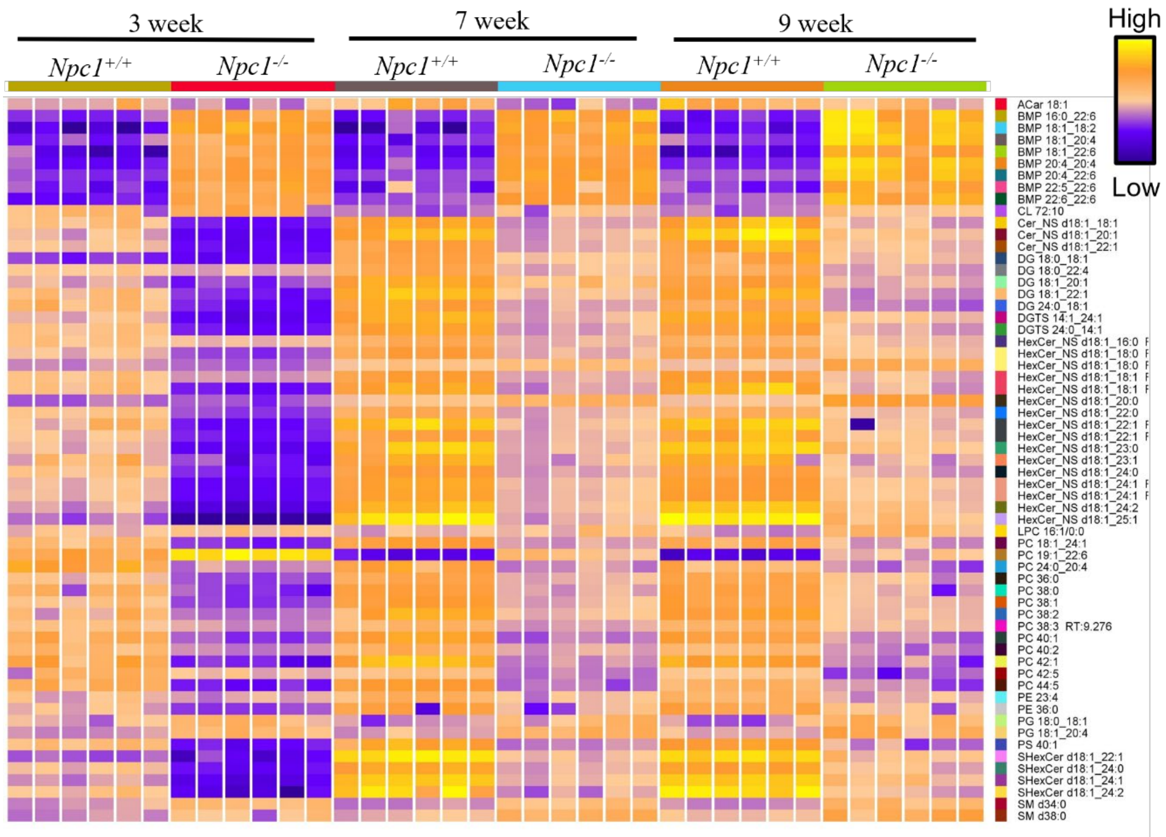


**Figure S6**. Heat map representing the lipids that are significantly different between 9-week *Npc1*$^{+/+}$ and *Npc1*$^{-/-}$ in the cortex. Unpaired t-tests were conducted between *Npc1*$^{-/-}$ and *Npc1*$^{+/+}$ for 9-week cortex positive ion mode data, and p-value < 0.05 and FC > 1.5 were considered significant. The color scale represents the normalized and baselined Log2 peak area of each lipid.

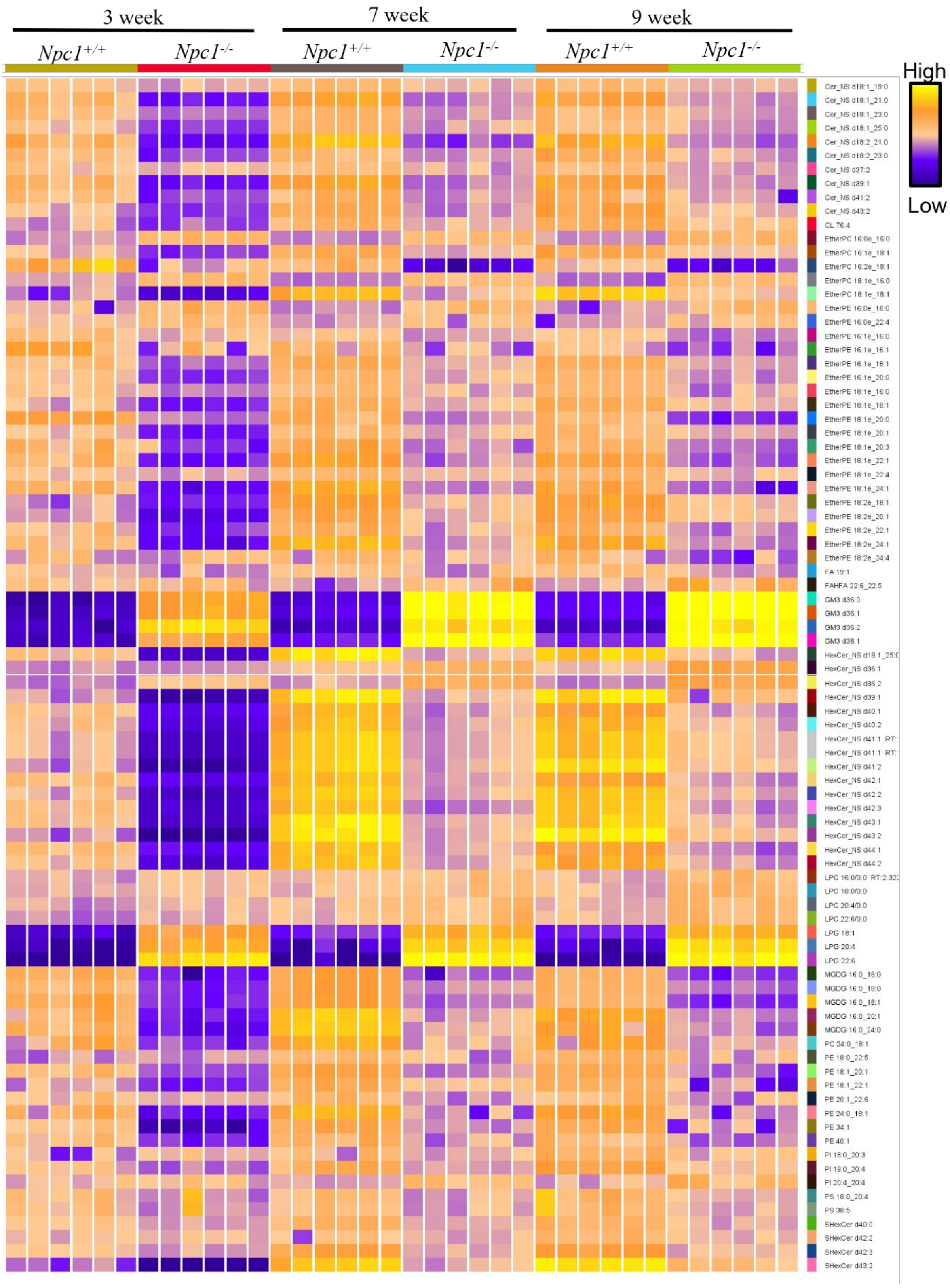


**Figure S7**. Heat map representing the lipids that are significantly different between 9-week *Npc1*$^{+/+}$ and *Npc1*$^{-/-}$ in the cortex. Unpaired t-tests were conducted between *Npc1*$^{-/-}$ and *Npc1*$^{+/+}$ for 9-week cortex negative ion mode data, and p-value < 0.05 and FC > 1.5 were considered significant. The color scale represents the normalized and baselined Log2 peak area of each lipid.

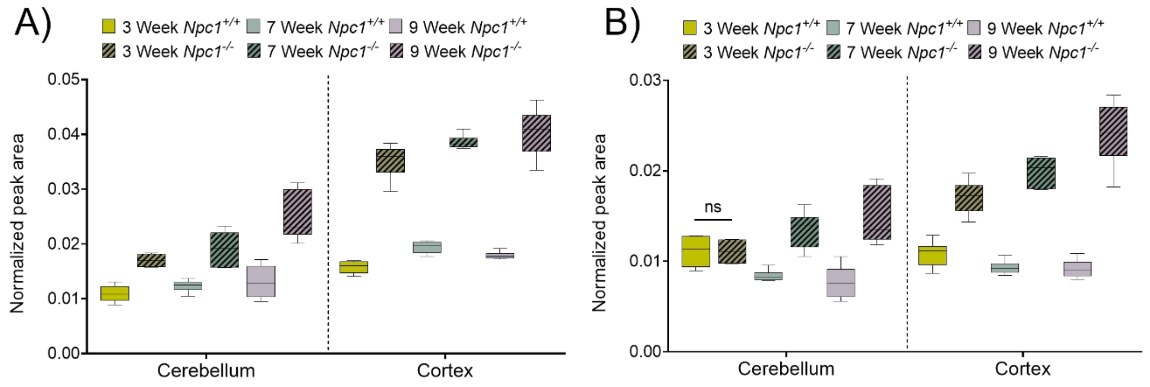


**Figure S**8. Increased levels of A) sphingosine and B) dihydro sphingosine (sphinganine) in the *Npc1*$^{-/-}$ cerebellum and cortex tissues. Peak areas normalized to the internal standards were used to create the box and whisker plots. An unpaired t-test was conducted to determine the p-value. p-value > 0.05 is considered as nonsignificant (ns).
